# A blunted GPR183/oxysterol axis during dysglycemia results in delayed recruitment of macrophages to the lung during *M. tuberculosis* infection

**DOI:** 10.1101/2022.01.05.475168

**Authors:** Minh Dao Ngo, Stacey Bartlett, Helle Bielefeldt-Ohmann, Cheng Xiang Foo, Roma Sinha, Buddhika Jayakody Arachige, Sarah Reed, Thomas Mandrup-Poulsen, Mette Marie Rosenkilde, Katharina Ronacher

**Author notes:** Correspondence: Katharina Ronacher.

## Abstract

We previously reported that the oxidised cholesterol-sensing receptor GPR183 is significantly downregulated in blood from tuberculosis (TB) patients with diabetes compared to TB patients without co-morbidities and that lower GPR183 expression in blood is associated with more severe pulmonary TB on chest-x-ray consistent with observations in dysglycemic mice. To further elucidate the role of this receptor and its endogenous high affinity agonist 7α,25-dihydroxycholesterol (7α,25-OHC) in the lung, we studied high fat diet (HFD)-induced dysglycemic mice infected with *M*.*tuberculosis*.

We found that the 7α,25-OHC-producing enzymes cholesterol 25-hydroxylase (CH25H) and cytochrome P450 family 7 subfamily member B1 (CYP7B1) were highly upregulated upon *M. tuberculosis* infection in the lungs of normoglycemic mice, and this was associated with increased expression of GPR183 indicative of effective recruitment of GPR183-expressing immune cells to the site of infection. We demonstrated that CYP7B1 was predominantly expressed by macrophages in the centre of TB granulomas. Expression of CYP7B1 was significantly blunted in lungs from HFD-fed dysglycemic animals and this coincided with delayed recruitment of macrophages to the lung during early infection and more severe lung pathology. GPR183 deficient mice similarly had reduced macrophage recruitment during early infection demonstrating a requirement of the GPR183/oxysterol axis for macrophage infiltration into the lung in TB.

Together our data demonstrate that oxidised cholesterols and GPR183 play an important role in positioning macrophages to the site of *M. tuberculosis* infection and that this is impaired by HFD-induced dysglycemia, adding a mechanistic explanation to the poorer TB outcomes in patients with diabetes.

## BACKGROUND

Type 2 diabetes (T2D) increases the risk for developing active tuberculosis (TB). TB patients with T2D co-morbidity are at higher risk of adverse TB treatment outcomes and increased mortality [1]. Several different animal models of diabetes show increased susceptibility to TB [2–8]. We previously reported that dysglycemic *M. tuberculosis* (Mtb)-infected mice had more severe TB with a trend towards higher lung bacterial burden, significantly lower pulmonary concentrations of tumor necrosis factor (TNF)-α and interferon (IFN)-γ during early infection (3 weeks post infection (p.i.)) accompanied by significantly worse lung pathology at 8 weeks p.i. compared to normoglycemic control animals [2]. While various mechanisms likely contribute to an impaired host defense in subjects with hyperglycemia, chronic low-grade inflammation leading to defects in the innate immune responses against Mtb and the subsequent delay in activating adaptive immune responses have been suggested as cellular mechanisms of TB susceptibility [6]. This arises, at least in part, through impaired recognition of Mtb by diabetic alveolar macrophages (AMs) [8].

After infection of AMs with Mtb, the lungs are infiltrated by various cell populations, with neutrophils, interstitial monocytes, macrophages, dendritic cells [9], and eosinophils [10] being recruited to the lung during the first two weeks of infection. This cellular migration requires effective chemotactic signals to direct the immune cells to the lung, and signals from Mtb-infected AMs likely to play a critical part [9].

In addition to classical chemokines, oxidised cholesterols so called oxysterols, have been identified as chemoattractants controlling the movement of distinct immune cells to position them to specific tissues [11]. The oxysterol 7α,25-OHC, the most potent endogenous agonist for the G-protein coupled receptor GPR183, is produced from cholesterol via two hydroxylation steps by the enzymes Cholesterol 25-hydroxylase (CH25H) and Cytochrome P450 Family Subfamily B Member 1 (CYP7B1), respectively. This oxysterol can subsequently be metabolised by hydroxyl D-5-steroid dehydrogenase, 3β- and steroid D-isomerase 7 (HSD3B7) into 4-cholesten-7α,25-ol-3-one [12]. Besides B cells and T cells, GPR183 is also expressed on innate immune cells such as macrophages and natural killer cells [13, 14]. GPR183 and 7α,25-OHC are important for chemotactic distribution of immune cells to secondary lymphoid organs [15–21] and positioning of innate lymphoid cells (ILCs) in the gut [22, 23].

However, only few studies evaluated the role of oxysterols in the lung.Various oxysterols were increased in bronchoalveolar lavage fluid from asthma patients and associated with infiltration of eosinophils, neutrophils and lymphocytes to the lung [24]. The upregulation of CH25H and CYP7B1 has also been demonstrated in the inflammated lungs of patients with chronic obstructive pulmonary disease [25]. In a murine model of lipopolysaccharide (LPS)-induced acute lung inflammation 25-hydroxycholesterol (25-OHC) was upregulated in the lung [26]. However, the role of oxysterols and GPR183 in infectious respiratory diseases including TB has not been investigated. We previously observed significantly lower GPR183 expression in blood from TB patients with T2D compared to TB patients without co-morbidities and, low GPR183 expression correlated with increased TB disease severity assessed by chest x-ray [27], suggesting GPR183 and 7α,25-OHC may be important in TB pathogenesis.

In the present study, we investigated the role of GPR183 and 7α,25-OHC in immune cell recruitment to the Mtb-infected lung in normoglycemic and dysglycemic mice. We report that CH25H and CYP7B1 are upregulated in the lung in response to Mtb infection. In dysglycemic mice, however, the Mtb-infection induced induction of CYP7B1 expression was absent [28] and associated with reduced macrophage infiltration into the lung. Similarly mice genetically deficient for GPR183, who have higher Mtb loads during early infection [27], had reduced macrophage recruitment upon Mtb infection suggesting a requirement of GPR183 for effective macrophage infiltration into the lung and effective containment of Mtb.

## METHODS

### Murine Models and Mtb infection

We have previously characterised the high fat diet (HFD) murine model of dysglycemia and tuberculosis [2]. Briefly, six-week-old male C57BL/6 mice were fed a lard based HFD (HFD), which contained 43% available energy as fat (total fat: 23.50%, SF04-001, Specialty Feeds, Western Australia). Control animals were fed normal chow diet (NCD) with 12% available energy from fat for the same period (total fat: 4.60%, Standard rodent diet, Specialty Feeds, Western Australia). At 12 weeks on the respective diets mice were infected with approximately 150 cfu of Mtb H_37_R_v_ as previously described [2]. GPR183 KO mice were infected with Mtb as previously described [27]. Tissues were collected for downstream analysis at either 2, 3, 5 or 8 weeks p.i. as indicated in figure legends.

### RNA Extraction and qRT-PCR

RNA was isolated from lung and blood using Isolate II RNA mini kit protocol (Bioline Reagents Ltd., London, UK) with slight modification. Briefly, blood cell pellet and lung lobes were homogenized in Trizol, vigorously mixed with chloroform (2.5:1) and centrifuged at 12,000 x g for 15 min at 4°C. The RNA in the aqueous phase was precipitated by mixing in cold 70% ethanol (1:2.5) followed by column-based RNA isolation using kit protocol including DNase treatment to remove genomic DNA contamination. Complementary DNA was synthesized using 2 µg of RNA and the Tetro cDNA synthesis kit (Bioline Reagents Ltd., London, UK) according to manufacturer’s instructions. Gene expression analysis was performed by quantitative real time PCR (qRT-PCR) with SensiFAST™ SYBR® Lo-ROX Kit (Bioline Reagents Ltd., London, UK) run on the QuantStudio™ 7 Flex Real-Time PCR System (Applied Biosystems). All gene expression levels were normalized to Hprt1 internal controls in each sample, and the fold changes were calculated using the 2^−ΔΔCT^ method. The list of primers used is given in Table S1.

### Immunohistochemistry

Formalin-fixed paraffin-embedded (FFPE) lung sections were dewaxed in xylene before hydrating with decreasing ethanol changes. Endogenous peroxidase activity was blocked with 3% hydrogen peroxide for 10 min and antigen retrieved using proteinase K (Sigma-Aldrich, P6556) in Tris-EDTA pH8 buffer for 4 min at room temp. Non-specific antibody binding was blocked using Background Sniper (Biocare Medical, Concord, CA). Immuno-labeling was performed with rabbit antibodies against Iba1 (Novachem, 019-19741), CYP7B1 (BIOSS BS-5052R), or isotype control (Rabbit IgG, ThermoFisher 31235) diluted in Da Vinci Green Diluent (Biocare Medical) for 120 min. Sections were subsequently incubated with HRP-conjugated goat anti-rabbit (Abcam, ab6721). To develop reactions, diaminobenzidine (Dako, Agilent) was used as per the manufacturer’s instructions for 2 min. Sections were counterstained with Mayer’s hematoxylin (Sigma-Aldrich), imaged using VS120 slide scanner (Olympus, TYO, JP), and analyzed using Visiopharm® software (Visiopharm, DK) or a Olympus BX50 microscope via Olympus CellSens standard software 7.1 (Olympus).

### Mass spectrometric quantitation of 25-OHC and 7α,25-OHC in lung homogenates

The oxysterol quantification method was adapted from McDonald et al.[29]. Briefly, Mtb-infected lung homogenates were treated with formalin for 24 h before removing samples out of PC3 lab, following an approved safety protocol for Mtb infected samples. Oxysterols were extracted using a 1:1 dichloromethane:methanol solution containing 50 µg/mL mL butylated hydroxytoluene (BHT) in a 30°C ultrasonic bath. Tubes were flushed with N_2_ to displace oxygen, sealed with a polytetrafluoroethylene (PTFE)-lined screw cap, and incubated at 30°C in the ultrasonic bath for 10 min. Following centrifugation (3,500 rpm, 5 min, 25°C), the supernatant from each sample was decanted into a new tube. For liquid-liquid extraction, Dulbecco’s phosphate-buffered saline (DPBS) was added to the supernatant, agitated and centrifuged at 3,500 rpm for 5 min at 25°C. The organic layer was recovered and evaporated under N_2_ using a 27-port drying manifold (Pierce; Fisher Scientific, Fair Lawn, NJ). Oxysterols were isolated by solid-phase extraction (SPE) using 200 mg, 3 mL aminopropyl SPE columns (Biotage; Charlotte, NC). The samples were dissolved in 1 ml of hexane and transferred to the SPE column, followed by a rinse with 1 ml of hexane to elute nonpolar compounds. Oxysterols were eluted from the column with 4.5 ml of a 23:1 mixture of chloroform:methanol and dried under N_2_. Samples were resuspended in 100 µl of warm (37°C) 90% methanol, 0.1% DMSO, and placed in an ultrasonic bath for 5 min at 30°C. A standard curve was extracted for 25-OHC (Sigma-Aldrich, H1015) and 7α,25-OHC (Sigma-Aldrich, SML0541) using the above method.

Samples were analysed on AB Sciex QTRAP® 5500 (ABSCIEX, Redwood City, CA) mass spectrometer coupled to a Shimadzu Nexera2 UHPLC. A Kinetex Pentafluorophenyl (PFP) column (50 × 2.1mm, 1.7µM, Phenomenex) was used for the separation of 25-OHC and 7α,25-OHC from other oxysterols. Mobile phase used for separation were, A - 0.1% formic acid with water and B - 100% acetonitrile with 0.1% formic acid. Five µL of sample were loaded at 0.4mL/min and separated using linear gradient with increasing percentage of acetonitrile. Samples were washed for 1.3 min after loading with 40% mobile phase B followed by linear gradient of 40%-99% over 6min. The column was washed with 99% mobile phase B for 2 min followed by equilibration with 40% B 2 min before next injection. Column oven and auto-sampler were operated 60°C and 6°C, respectively. Elution of analytes from the column was monitored in positive ion mode (ESI) with multiple reaction monitoring on ABSciex QTRAP mass spectrometer equipped with Duospray ion source, which was operated at temp 550°C, ionspray voltage of 5500, curtain gas (CUR) of 30, ion source gas1 (GS1) of 65 and ion source gas 2 (GS2) of 50. Quadrupole 1 and 3 were operated at unit mass resolution at all time during the experiment. MRM pairs 385.3 > 367.3, 385 >133, 385.3 > 147.1 and 367 >147.1 were monitored for 25-OHC and for 7α,25-OHC following MRM pairs were used 436.3>383.3, 383.2 > 365.3, 383.2 > 147.3, 383.2 > 159.0. Deuterated 25-OHC (Sapphire Bioscience, 11099) used as an internal standard. De-clustering potential (DP), Collision energy (CE), Entrance (EP) and collision cell exit potential (CXP) were optimised for each MRM pair to maximise the sensitivity. Data processed using AbSciex MultiQuant software (Version 3.0.3)

### Ethics Statement

All experiments were carried out in accordance with protocols approved by the Health Sciences Animal Ethics Committee of The University of Queensland (MRI-UQ/413/17) and performed in accordance with the Australian Code of Practice for the Care and Use of Animals for Scientific Purposes.

## RESULTS

### Mtb infection increases CH25H and CYP7B1 expression in the lung which results in increased oxysterol production

To investigate whether Mtb-infection induces the production of oxidised cholesterols in the lung we infected mice with Mtb and determined the mRNA expression of oxysterol producing enzymes. Cholesterol is hydroxylated by the enzyme CH25H to form 25-OHC and subsequently hydroxylated by CYP7B1 to form 7α,25-OHC (Figure 1A), the high affinity agonist for GPR183. 7α,25-OHC can be further metabolized by the enzyme HSD3B7. We found that Mtb infection significantly upregulated the expression of both *Ch25h* (Figure 1B, black bars) and *Cyp7b1* (Figure 1C, black bars), while *Hsd3b7* was downregulated in the lung at 3 weeks after infection (Figure 1D, black bars). To determine whether increased expression of these enzymes results in increased oxysterol production we performed mass spectrometry on lung homogenates for detection of 25-OHC and 7α,25-OHC. Mtb infection significantly increased 25-OHC production compared to 25-OHC concentrations in uninfected lung homogenates (Figure 1E, black bar vs. dotted line). We were unable to accurately determine 7α,25-OHC concentrations in the lung homogenates as they were below the detection limits of the system. However, these results demonstrate that Mtb infection induces expression of *Ch25h* and *Cyp7b1* likely via interferons [30] and results in increased production of 25-OHC and likely also increased production of 7α,25-OHC, both being ligands for GPR183.

**Figure 1:**
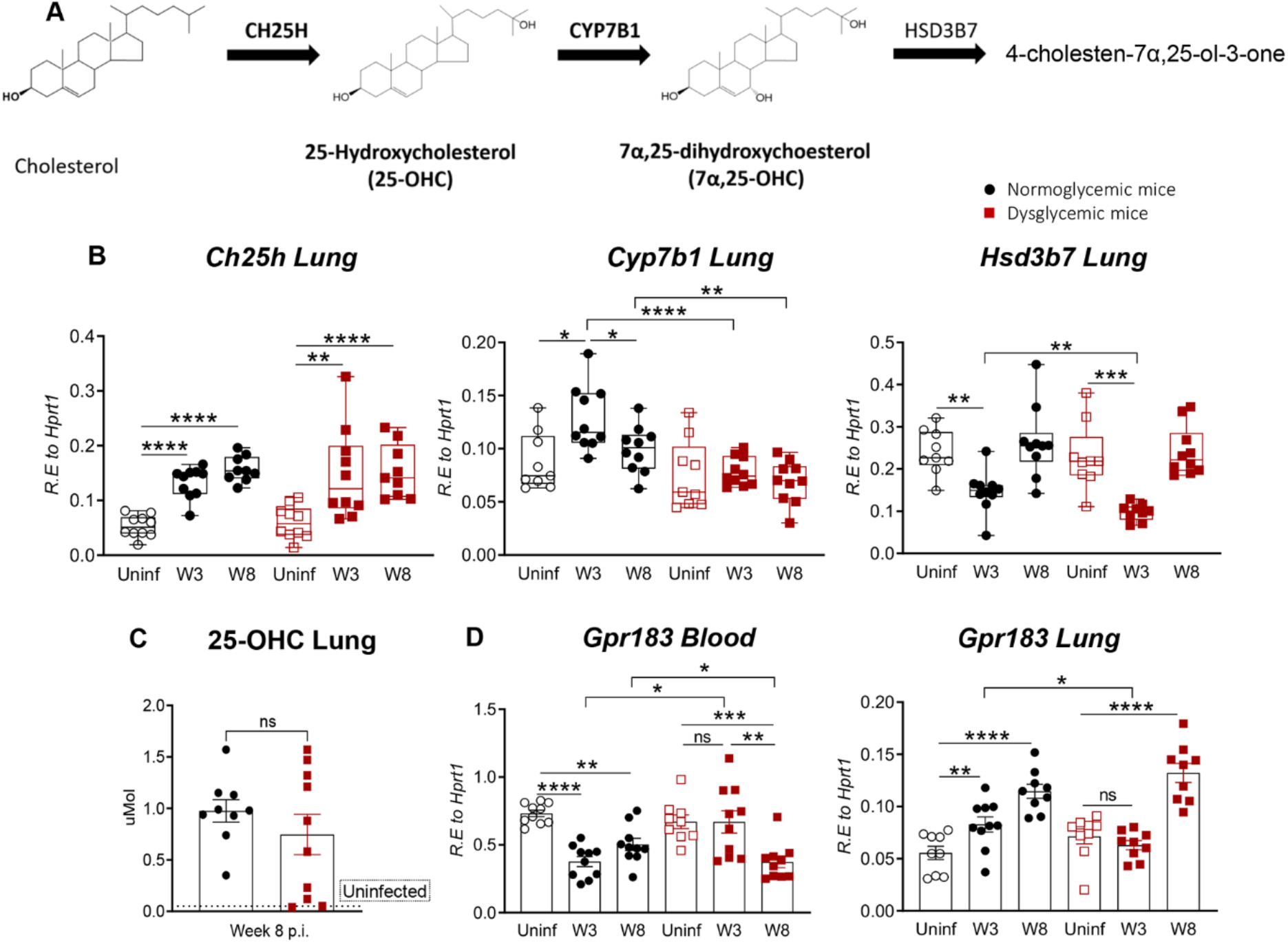
Expression of *Ch25h, CYP7B1, GPR183* and 25-OHC concentrations in lungs from uninfected and Mtb-infected normoglycemic and dysglycemic mice. **(A)** The biosynthetic pathway of 25-OHC and 7α,25-OHC. Normoglycemic and dysglycemic mice were infected with ∼100 CFUs aerosolized Mtb H_37_R_v_, and mRNA expression of **(B)** *Ch25h, Cyp7b1* and *Hsd3B7* was measured by qRT-PCR at 3-and-8 weeks post infection. (**C**) Concentrations of 25-OHC were measured in the lungs 8 weeks post infection. Uninfected mice fed a HFD or NCD diet are represented by the dotted line. Expression of (**D**) blood and lung *Gpr183* mRNA at 3 and 8 weeks post infection was determined by qRT-PCR. Data are means ± SEM of n=9-10 infected mice/group analyzed from one experiment. Circles represent normoglycemic mice and squares represent dysglycemic mice. Data analysis was performed by Mann-Whitney U test. ns = not significant, *p < 0.05, **p<0.01, ***p < 0.001 and ****p < 0.0001.

### Oxysterol production is associated with increased GPR183 expression

We have previously reported that lower *GPR183* expression in blood from TB patients is associated with increased TB disease severity on chest x-ray [27] and hypothesized that this is due to chemoattraction of GPR183-expressing immune cells towards a gradient of 25-OHC and 7α,25-OHC to the site of disease, the lung. Consistent with our observation in humans we found that in Mtb-infected mice *Gpr183* expression decreased in blood compared to uninfected animals (Figure 1 F, black bars), while *Gpr183* expression in lung significantly increased upon Mtb infection at both week 3 and 8 p.i. (Figure 1G, black bars). These data suggest that oxysterol sensing GPR183-expressing immune cells migrate towards the lung upon infection.

### Dysglycemia blunts Mtb-induced expression of CYP7B1 and GPR183 in the lung

Since diabetes is a well known risk factor for TB and we previously showed that mice with HFD-induced dysglycemia have more severe TB [2], we next investigated whether this is linked to changes in oxysterol production in the lung. We observed that Mtb-infection in dysglycemic animals induced *Ch25h* expression similar to in normoglycemic animals (Figure 1B red vs. black bars), and the concentrations of 25-OHC were similarly comparable between the animals (Figure 1E red vs. black bar). Interestingly, however, the expression of *Cyp7b1* was significantly blunted by dysglycemia and was consistently lower in dysglycemic compared to normoglycemic mice both at week 3 and week 8 p.i. (Figure 1C, red vs. black bars). This suggests that the production of 7α,25-OHC is impaired during dysglycemia likely due to insulin resistance[28]. Consistent with this, in dysglycemic animals we did not oberve a rapid decrease of *Gpr183* in blood (Figure 1 F) or a rapid increase of *Gpr183* in lung (Figure 1G) within the first three weeks post infection such as observed in normoglycemic animals. *Gpr183* increased only at week 8 p.i. in the lungs of HFD-fed animals.

Taken together, these results demonstrate that Mtb infection results in the production of oxysterols which facilitate the rapid migration of GPR183-expressing immune cells from the periphery to the lung. This GPR183-oxysterol axis isblunted during dysglycemia resulting in delayed recruitment of immune cells to the Mtb-infected lung.

### Dysglycemia blunts CYP7B1 protein in the lung after Mtb infection

To further confirm whether the reduced mRNA expression of *Cyp7b1* in dysglycemic animals translates into lower protein expression of CYP7B1 we performed immunohistochemical labeling of mouse lung sections with a CYP7B1 specific antibody. We observed that positive signals of CYP7B1 started to accumulate around blood vessels and bronchioles by 3 weeks p.i., with intense signals of CYP7B1 found mostly located in the center of granulomas by week 8 p.i. (Figure 2A). Quantification of positive immunolabeling confirmed the upregulation of CYP7B1 in the lung post infection (Figure 2B), and percent area of CYP7B1(+) was significantly reduced in dysglycemic mice compared to normoglycemic controls at 8 week p.i. (Figure 2B). These results suggest that distinct cell populations involved in granuloma formation drive CYP7B1 expression.

**Figure 2:**
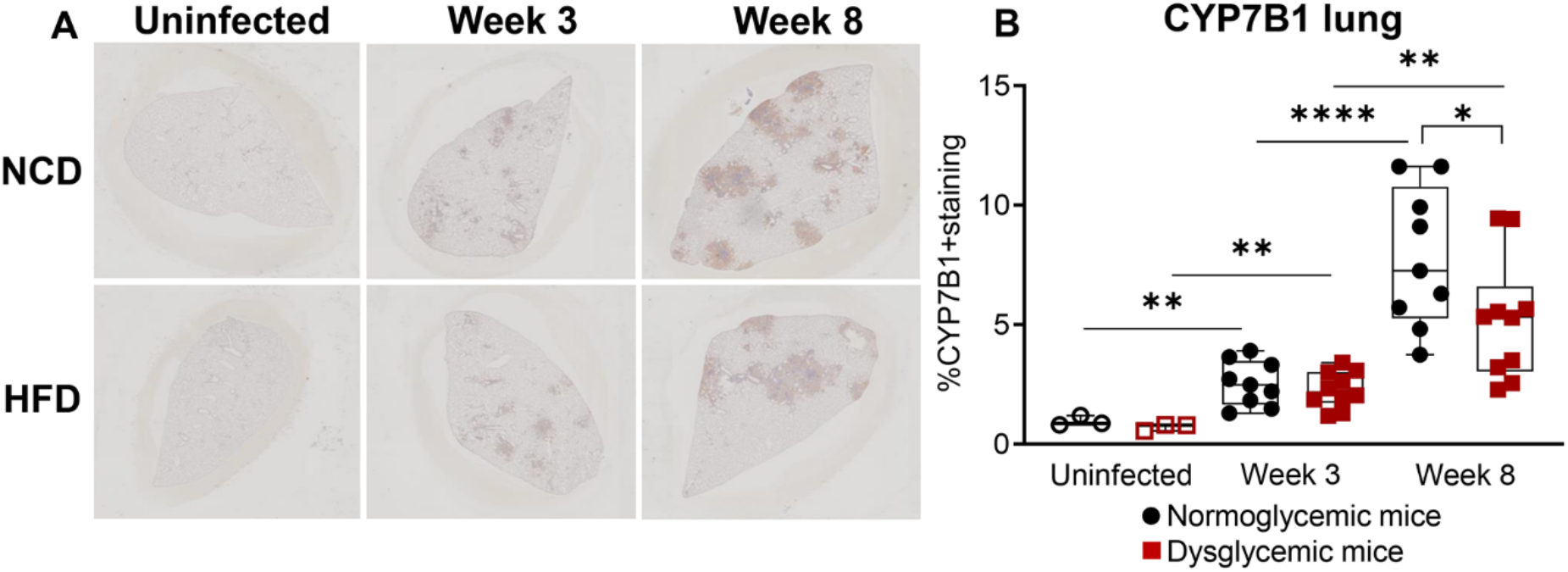
Mice with dysglycemia have lower CYP7B1 at protein levels compared to control normoglycemic mice. **(A)** Representative images of immunohistochemical labeling of CYP7B1 in lung sections from dysglycemic vs. normoglycemic mice either uninfected or 3 weeks and 8 week after Mtb infection. **(B)** Quantitative analysis of CYP7B1 immunolabeled areas. Data are means ± SEM of n=9-10 infected mice/group analyzed from one experiment. Circles represent normoglycemic mice and squares represent dysglycemic mice. Data analysis was performed by Mann-Whitney U test. *p < 0.05.**p < 0.01 and ****p < 0.0001.

### CYP7B1 is expressed by alveolar macrophages and infiltrating macrophages upon Mtb infection

We next investigate which cell type in the lung produces CYP7B1 upon Mtb infection and found that CYP7B1 was most abundant in alveolar macrophages, indentified as large round cells with unsegmented nuclei inside alveolar spaces, of Mtb-infected but not in uninfected animals (Figure 3A, middle low image). At week 8 p.i., intense signals of CYP7B1 were found in the center of granulomas (Figure 3A, right low image). We confirmed that CYP7B1 expression was macrophage derived by immunolabeling lung sections with the macrophage-specific marker ionized calcium binding adaptor molecule 1 (Iba1), the distribution of which overlapped with CYP7B1 expression (Figure 3B). Cells positive for the Iba1 signal started to appear around blood vessels and bronchioles by week 3 p.i., indicating that Iba1^+^ macrophages from circulation infiltrated into the lung after Mtb infection and these Iba1^+^ macrophages expressed CYP7B1.

**Figure 3:**
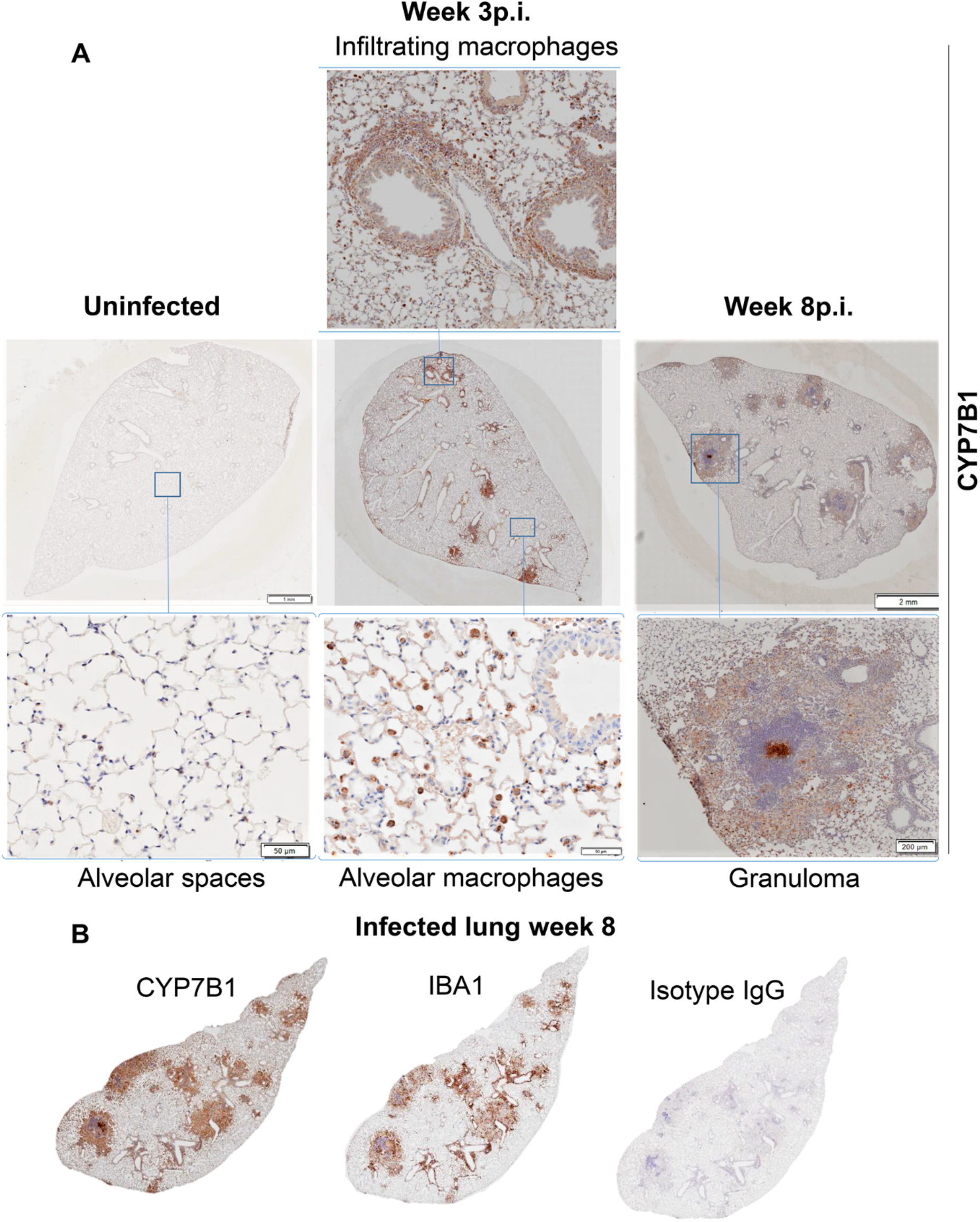
CYP7B1 protein expression in alveolar macrophages and in the center of TB granulomas. **(A)** Representative images of CYP7B1 immunolabeling on lung sections from uninfected and Mtb-infected animals at 3 and 8 weeks p.i..(**B**) Representative image of serial sections immunolabeled for Iba1 from the lung of mice 8 weeks post infection.

Taken together, we demonstrate that CYP7B1 is upregulated in both resident alveolar macrophages and infiltrating macrophages upon Mtb infection and is highly expressed in the center of granulomas. It is thus possible that the oxysterol/GPR183 axis plays an important role in positioning of leukocytes around Mtb infected macrophages in the TB granuloma.

### Dysglycemia leads to lower macrophage infiltration to the Mtb-infected lung

We next assessed whether dysglycemia impacts macrophage migration into the Mtb-infected lung. We found that there was more than a five-fold increase in macrophages within the first three weeks p.i. in normoglycemic animals; however, macrophage recruitment was significantly lower in dysglycemic mice at that timepoint (Figure 4A, B). Together these results suggest that the lower *Gpr183* expression we observed in dysglycemic animals is due to a reduction in macrophage infiltration to the lung during early Mtb infection. We next postulated that the impaired migration of macrophages to the Mtb infected lung is oxysterol/GPR183 dependent.

**Figure 4:**
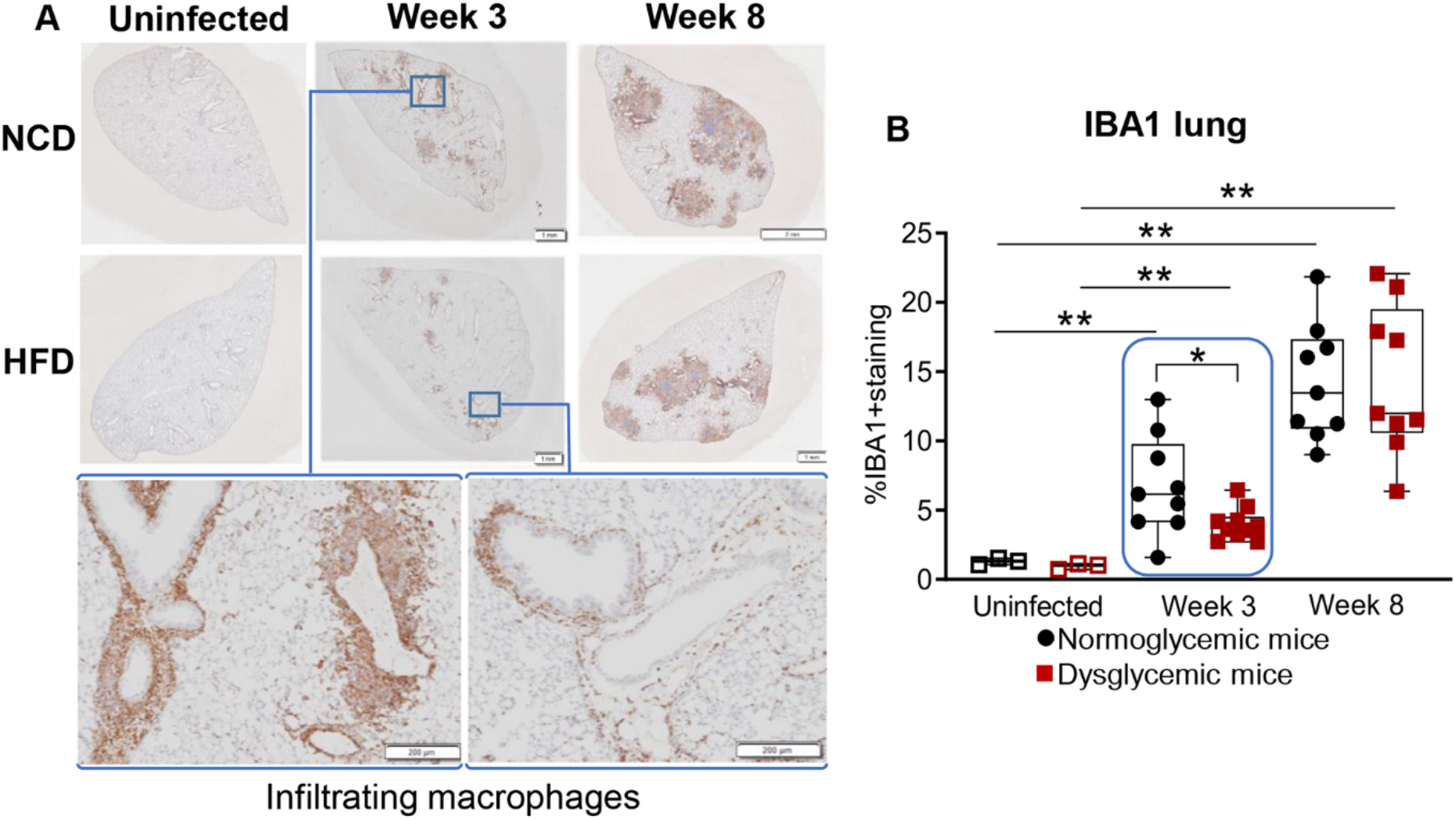
Reduced infiltration of macrophages in mice with dysglycemia at 3 weeks p.i.. **(A)** Representative images of the macrophage marker Iba1 immunolabeled lung sections from uninfected and Mtb-infected animals at 3-and-8 weeks p.i. **(B)** Quantitative analysis of Iba1(+) labeled areas. Data are means ± SEM of n=9-10 mice from infected groups analyzed from one experiment. Circles represent normoglycemic mice and squares represent dysglycemic mice. Data analysis was performed by Mann-Whitney U test. *p < 0.05 and **p < 0.01.

### GPR183 is required for efficient macrophage infiltration to the lung during early Mtb infection

To investigate whether the GPR183/7α,25-OHC axis is required for macrophage infiltration into the Mtb infected lung, we performed experiments in GPR183KO mice. Previously, we reported that GPR183KO mice presented with increased Mtb burden compared to WT animals at 2 weeks after infection with Mtb H_37_R_v_, an effect that disappeared at 5 weeks post infection [27]. This suggests that GPR183 plays an important role during the early innate immune response to Mtb infection. We speculated that absence of GPR183 could alter the recruitment and distribution of immune cells to the lung in the context of TB disease. We found that pulmonary macrophage infiltration was significantly reduced in GPR183KO compared to WT mice at 2 weeks p.i.. However, the absence of GPR183 does not result in prolonged macrophage deficiency in the lungs in infected mice as by week 5 p.i, significantly more macrophages infiltrate the lungs of GPR183KO mice vs WT mice (Figure 5) indicative of a compensatory GPR183/7α,25-OHC independent mechanism of macrophage recruitment to the lung at that later timepoint during infection.

**Figure 5:**
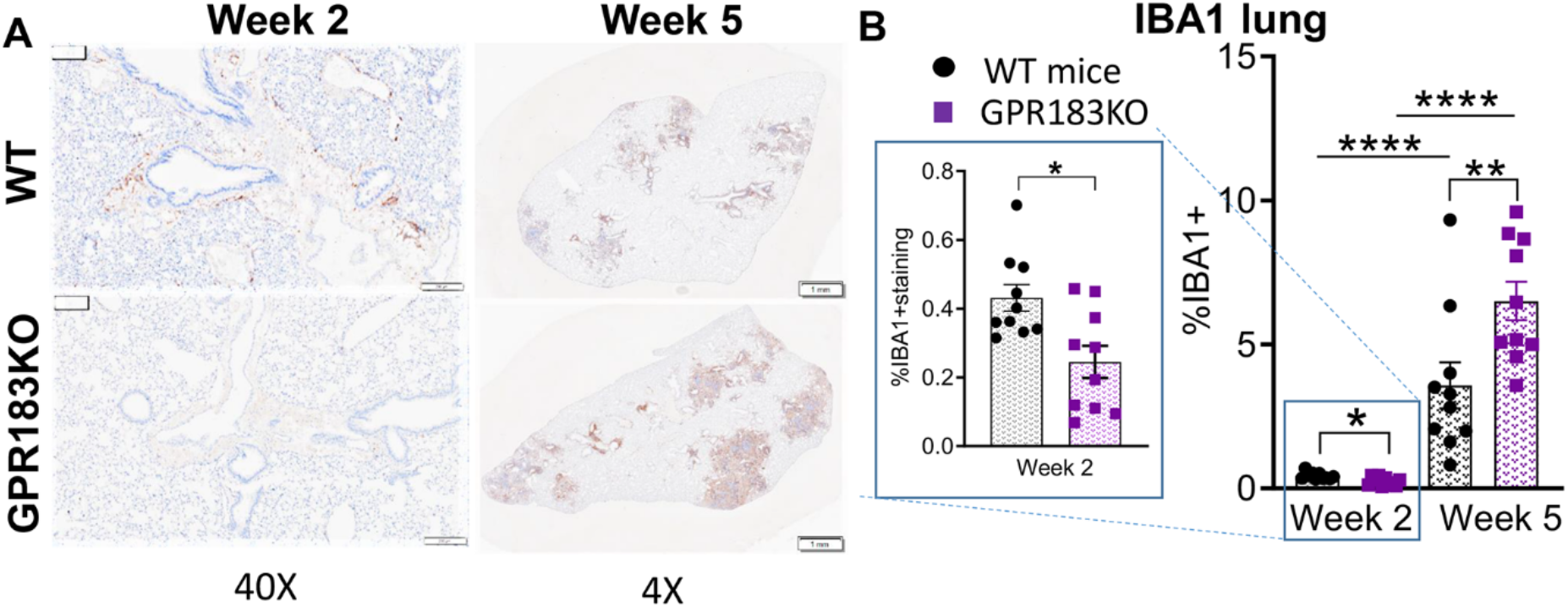
Reduced macrophages infiltration to the lung of GPR183KO mice at 2 weeks after Mtb infection. WT and GPR183KO mice were infected with Mtb as previously described [27]. **(A)**. Representative Iba1 immunolabeled lung sections from Mtb-infected WT and GPR183KO mice at 2 and 5 weeks p.i. **(B)**. Quantification of Iba1+ labeled areas. Data are means ± SEM of n=10 infected mice/group analyzed from one experiment. Circles represent C57/BL6 wildtype mice and squares represent GPR183KO mice. Data analysis was performed by Mann-Whitney U test. *p < 0.05; **p<0.01 and ****p < 0.0001.

These results indicate that GPR183 is necessary for effective recruitment of inflammatory and anti-microbial macrophages to the lung during the early Mtb infection. This early impairment of macrophage migration to the lung likely contributed to higher lung bacterial numbers in GPR183KO [27] and dysglycemic mice [2].

## DISCUSSION

In this study we demonstrated a role for oxysterols and GPR183 in positioning of immune cells to the Mtb-infected lung with potential implications for other bacterial and viral respiratory tract infections. Previous studies have illustrated the importance of GPR183 in migration of immune cells to secondary lymphoid organs including the positioning of B cells in lymphoid tissues [16, 17], dendritic cells to the marginal zone bridging channels in the spleen [16, 31], T cells in T cells zone or more recently, the localization of ILC3 to lymphoid structures in the colon [22, 32]. Yet, the role of oxysterols and GPR183 in positioning of immune cells in the lung is largely unexplored. Jia et al. showed in a mouse model of COPD that GPR183 is required for the formation of inducible bronchus-associated lymphoid tissue (iBALT), a secondary lymphoid-like structure within the lung [25]. iBALTs are also formed during Mtb infection and correlate with protection [33]. The initial host determinants that govern the induction of iBALT formation during Mtb infection remain to be elucidated, but it is possible that oxysterols and GPR183 play a major role in iBALT formation during TB by serving as recruitment and retention signals. Our observation that GPR183 KO mice are more susceptible to Mtb infection during the first two weeks could at least in part be due to deficiencies in iBALT formation. However it is also possible that the increased Mtb burden is linked to the lower macrophage infiltration we observed during early infection in GPR183 KO mice [27]. A delayed innate immune activation results in delayed adaptive immune priming. Consistent with this GPR183KO mice demonstrated delays in *ex vivo* pro-inflammatory cytokine responses with significantly lower IFN-β, IFN-γ and a trend toward lower IL-1β production compared to WT controls [27]. We showed that during the later stages of infection, GPR183KO mice accumulated significantly more macrophages in the lung compared to WT animals through GPR183 independent mechanisms. These GPR183 independent mechanisms could include multiple other cellular targets of oxysterols involved in immune regulation [34] or could be chemokine mediated.

We found that a lack of GPR183 impacts mainly macrophage infiltration, even though this receptor is also expressed on other immune subsets. Consistent with our finding others demonstrated that T cells do not require GPR183 for migration into the Mtb-infected lung [35]. We showed that CYP7B1 is almost exclusively expressed in alveolar macrophages and infiltrating macrophages upon Mtb infection and not present in uninfected lungs. Thus, we identified macrophages as the major source of oxysterols in the Mtb-infected lung. Previous studies have reported high expression of CH25H by macrophages [13, 36–38] and we found that both CH25H and CYP7B1 are upregulated upon Mtb infection in both primary human monocytes and in THP-1 macrophages (data not shown) and this is likely mediated by interferons as CH25H is interferon inducible[30]. In a cigarette smoke-exposed mouse model, Jia et al demonstrated high expression of *Ch25h* in the airway epithelial cells [25], while Madenspacher et al. found that *Ch25h* and its product 25-OHC are highly expressed in resident lung alveolar macrophages, but not in macrophages from other compartments in LPS-exposed C57BL/6 mice [39]. In lymphoid tissues stromal cells have been reported to be the main *Ch25h* and *Cyp7b1* expressing cells and major contributors to 7α,25-OHC generation [31]. These data suggest that the cellular source of 7α,25-OHC producing enzymes varies dependent on the type of stimulus and local environment of the involved organs.

At the later timepoint 8 week p.i., CYP7B1 was abundantly expressed in the center of granulomas compared to surrounding regions. This suggests that 7α,25−OHC is highly produced at the center of granulomas to attract other macrophages and lymphocytes making the oxysterol/GPR183 an important element in positioning immune cells in the granuloma. Another interesting observation in our study is that the intensity of CYP7B1 signals is not uniform throughout all granulomas from the same lung lobe and likely reflects heterogeneity of developmental stages of granulomas in the Mtb-infected lung [40–42].

The concentrations of oxysterols in serum are modified by diabetes and obesity with some up-and some down-regulated [43, 44]. Aberrant oxysterol metabolism in diabetes is also present in the liver [45]. In an insulin resistant HFD-based mouse model chronic suppression of CYP7B1 was observed in the liver accompanied by reduced production of 25-OHC [28]. However whether oxysterol concentrations vary in the lung upon HFD feeding or diabetes has not been investigated. Here, we demonstrate the Mtb-induced upregulation of CYP7B1 is blunted during dysglycemia. Reduced CYP7B1 expression likely results in reduced 7α,25−OHC production. A limitation of our study was that we were unable to measure this oxysterol due to technical constraints. However a reduced 7α,25−OHC production in dysglycemic compared to normoglycemic mice can explain the delayed recruitment of macrophages to the site of infection. Delayed infiltration of myeloid cells to the Mtb-infected lung in diabetic mice has also been shown by others [6], which the authors attributed to aberrant chemokine production at a time when the significance of oxysterols in immune cell migration was not yet considered.

In summary, we have shown, that Mtb infection results in increased expression of the oxysterol producing enzymes CH25H and CYP7B1, increased production of 25-OHC and likely also 7α,25−OHC in the lung (Fig. 6). Expression of CYP7B1 is blunted in dysglycemic animals likely due to insulin resistance and associated with reduced macrophage infiltration during early infection, which is also observed in GPR183KO mice. We therefore demonstrated that the oxysterol/GPR183 axis is important for immune cell positioning and possibly granuloma formation in TB. Further studies are required to assess how administration of oxysterols or GPR183 ligands modifies TB pathogenesis and outcomes and whether such compounds can be exploited for host-directed therapies.

**Figure 6:**
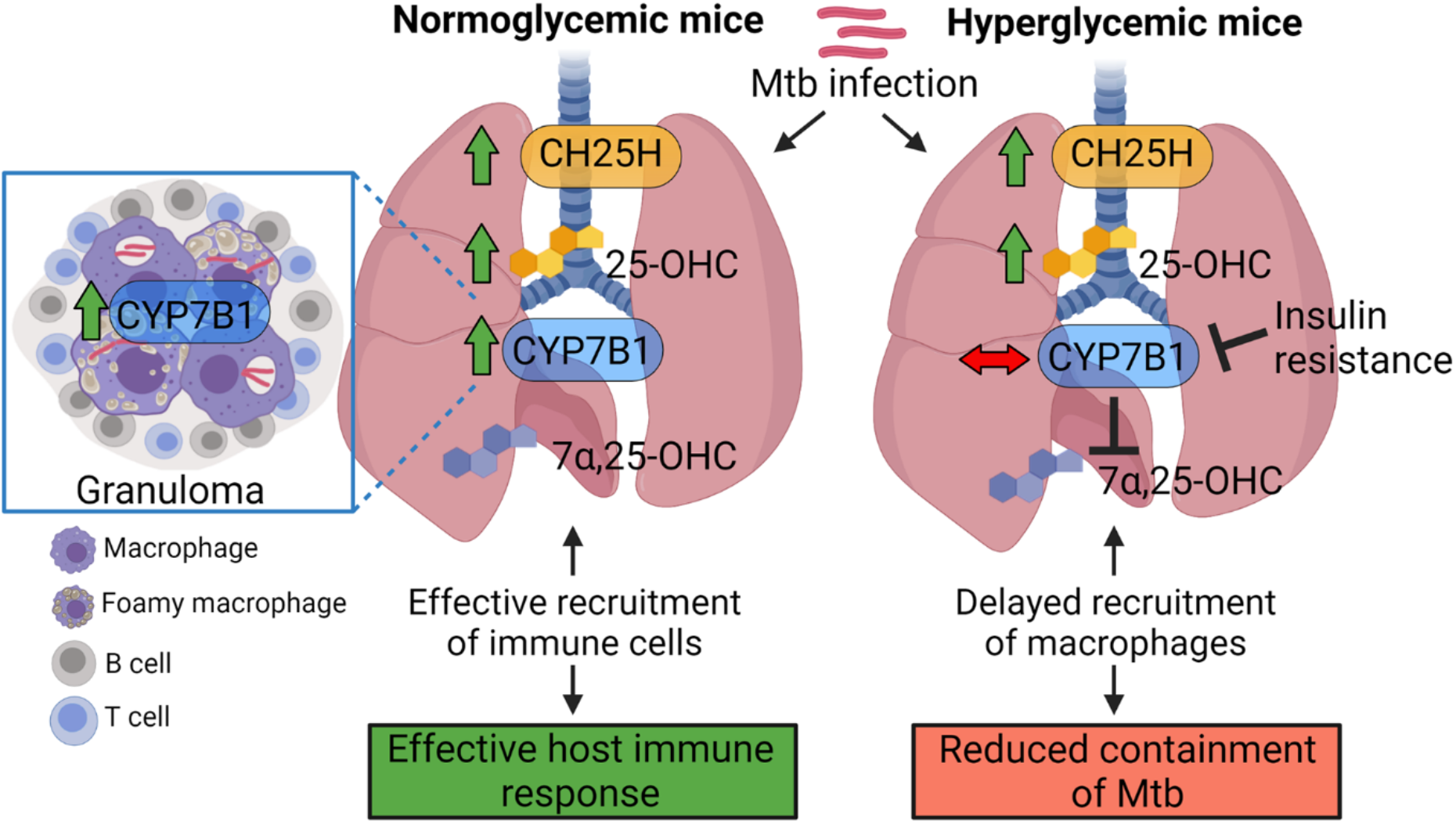
Schematic summary of the proposed role of the oxysterols 25-OHC and 7α,25-OHC and GPR183 in Mtb-infected normoglycemic and hyperglycemic mice. In the mouse lung after Mtb infection, the enzymes Cholesterol 25-hydroxylase (CH25H) and Cytpchrome P450 Family Subfamily B Member 1 (CYP7B1) are upregulated, along with the oxysterol 25-OHC and 7α,25-OHC resulting in effective recruitment of GPR183-expressing immune cells to the site of infection. CYP7B1 is predominantly expressed by macrophages in the centre of TB granuloma surrounded by a lymphocytes and therefore the oxysterol/GPR183 axis may contribute to positioning these immune cells in TB granulomas. In HFD-fed hyperglycemic mice, this pathway is altered, with CYP7B1 RNA and protein expression blunted in lungs coinciding with delayed recruitment of macrophages to the lung during early infection.

## Notes

### Competing Interest Statement

The authors have declared no competing interest.

## References

1. Critchley JA, Restrepo BI, Ronacher K, et al. Defining a Research Agenda to Address the Converging Epidemics of Tuberculosis and Diabetes: Part 1: Epidemiology and Clinical Management. Chest 2017; 152:165–73.

2. Sinha R, Ngo MD, Bartlett S, et al. Pre-Diabetes Increases Tuberculosis Disease Severity, While High Body Fat Without Impaired Glucose Tolerance Is Protective. Frontiers in Cellular and Infection Microbiology 2021; 11.

3. Yamashiro S, Kawakami K, Uezu K, et al. Lower expression of Th1-related cytokines and inducible nitric oxide synthase in mice with streptozotocin-induced diabetes mellitus infected with Mycobacterium tuberculosis. Clin Exp Immunol 2005; 139:57–64.

4. Martens GW, Arikan MC, Lee J, Ren F, Greiner D, Kornfeld H. Tuberculosis susceptibility of diabetic mice. Am J Respir Cell Mol Biol 2007; 37:518–24.

5. Sugawara I, Yamada H, Mizuno S. Pulmonary tuberculosis in spontaneously diabetic goto kakizaki rats. The Tohoku journal of experimental medicine 2004; 204:135–45.

6. Vallerskog T, Martens GW, Kornfeld H. Diabetic mice display a delayed adaptive immune response to Mycobacterium tuberculosis. Journal of immunology (Baltimore, Md : 1950) 2010; 184:6275–82.

7. Podell BK, Ackart DF, Obregon-Henao A, et al. Increased Severity of Tuberculosis in Guinea Pigs with Type 2 Diabetes: A Model of Diabetes-Tuberculosis Comorbidity. Am J Pathol 2014; 184:1104–18.

8. Martinez N, Ketheesan N, West K, Vallerskog T, Kornfeld H. Impaired Recognition of Mycobacterium tuberculosis by Alveolar Macrophages From Diabetic Mice. The Journal of infectious diseases 2016; 214:1629–37.

9. Mayer-Barber KD, Barber DL. Innate and Adaptive Cellular Immune Responses to Mycobacterium tuberculosis Infection. Cold Spring Harb Perspect Med 2015; 5:a018424.

10. Bohrer AC, Castro E, Hu Z, et al. Eosinophils are part of the granulocyte response in tuberculosis and promote host resistance in mice. Journal of Experimental Medicine 2021; 218.

11. Bah SY, Dickinson P, Forster T, Kampmann B, Ghazal P. Immune oxysterols: Role in mycobacterial infection and inflammation. The Journal of Steroid Biochemistry and Molecular Biology 2017; 169:152–63.

12. Mutemberezi V, Guillemot-Legris O, Muccioli GG. Oxysterols: From cholesterol metabolites to key mediators. Progress in Lipid Research 2016; 64:152–69.

13. Hannedouche S, Zhang J, Yi T, et al. Oxysterols direct immune cell migration via EBI2. Nature 2011; 475:524–7.

14. Rosenkilde MM, Benned-Jensen T, Andersen H, et al. Molecular Pharmacological Phenotyping of EBI2: AN ORPHAN SEVEN-TRANSMEMBRANE RECEPTOR WITH CONSTITUTIVE ACTIVITY*. Journal of Biological Chemistry 2006; 281:13199–208.

15. Preuss I, Ludwig M-G, Baumgarten B, et al. Transcriptional regulation and functional characterization of the oxysterol/EBI2 system in primary human macrophages. Biochem Biophys Res Commun 2014; 446:663–8.

16. Gatto D, Paus D, Basten A, Mackay CR, Brink R. Guidance of B Cells by the Orphan G Protein-Coupled Receptor EBI2 Shapes Humoral Immune Responses. Immunity 2009; 31:259–69.

17. Pereira JP, Kelly LM, Xu Y, Cyster JG. EBI2 mediates B cell segregation between the outer and centre follicle. Nature 2009; 460:1122–6.

18. Li J, Lu E, Yi T, Cyster JG. EBI2 augments Tfh cell fate by promoting interaction with IL-2-quenching dendritic cells. Nature 2016; 533:110–4.

19. Suan D, Nguyen A, Moran I, et al. T Follicular Helper Cells Have Distinct Modes of Migration and Molecular Signatures in Naive and Memory Immune Responses. Immunity 2015; 42:704–18.

20. Gatto D, Wood K, Caminschi I, et al. The chemotactic receptor EBI2 regulates the homeostasis, localization and immunological function of splenic dendritic cells. Nature Immunology 2013; 14:446–53.

21. Yi T, Cyster JG. EBI2-mediated bridging channel positioning supports splenic dendritic cell homeostasis and particulate antigen capture. eLife 2013; 2:e00757.

22. Chu C, Moriyama S, Li Z, et al. Anti-microbial Functions of Group 3 Innate Lymphoid Cells in Gut-Associated Lymphoid Tissues Are Regulated by G-Protein-Coupled Receptor 183. Cell Rep 2018; 23:3750–8.

23. Willinger T. Oxysterols in intestinal immunity and inflammation. Journal of Internal Medicine 2019; 285:367–80.

24. Shen ZJ, Hu J, Kashi VP, et al. Epstein-Barr Virus-induced Gene 2 Mediates Allergen-induced Leukocyte Migration into Airways. Am J Respir Crit Care Med 2017; 195:1576–85.

25. Jia J, Conlon TM, Sarker RS, et al. Cholesterol metabolism promotes B-cell positioning during immune pathogenesis of chronic obstructive pulmonary disease. EMBO Mol Med 2018; 10:e8349.

26. Bottemanne P, Paquot A, Ameraoui H, Guillemot-Legris O, Alhouayek M, Muccioli GG. 25-Hydroxycholesterol metabolism is altered by lung inflammation, and its local administration modulates lung inflammation in mice. The FASEB Journal 2021; 35:e21514.

27. Bartlett S, Gemiarto AT, Ngo MD, et al. GPR183 Regulates Interferons, Autophagy, and Bacterial Growth During Mycobacterium tuberculosis Infection and Is Associated With TB Disease Severity. Frontiers in Immunology 2020; 11.

28. Kakiyama G, Marques D, Martin R, et al. Insulin resistance dysregulates CYP7B1 leading to oxysterol accumulation: a pathway for NAFL to NASH transition. Journal of Lipid Research 2020; 61:1629–44.

29. McDonald JG, Smith DD, Stiles AR, Russell DW. A comprehensive method for extraction and quantitative analysis of sterols and secosteroids from human plasma. Journal of lipid research 2012; 53:1399–409.

30. Liu S-Y, Aliyari R, Chikere K, et al. Interferon-Inducible Cholesterol-25-Hydroxylase Broadly Inhibits Viral Entry by Production of 25-Hydroxycholesterol. Immunity 2013; 38:92–105.

31. Yi T, Wang X, Kelly Lisa M, et al. Oxysterol Gradient Generation by Lymphoid Stromal Cells Guides Activated B Cell Movement during Humoral Responses. Immunity 2012; 37:535–48.

32. Emgård J, Kammoun H, García-Cassani B, et al. Oxysterol Sensing through the Receptor GPR183 Promotes the Lymphoid-Tissue-Inducing Function of Innate Lymphoid Cells and Colonic Inflammation. Immunity 2018; 48:120–32.e8.

33. Dunlap MD, Prince OA, Rangel-Moreno J, et al. Formation of Lung Inducible Bronchus Associated Lymphoid Tissue Is Regulated by Mycobacterium tuberculosis Expressed Determinants. Frontiers in Immunology 2020; 11.

34. Reinmuth L, Hsiao C-C, Hamann J, Rosenkilde M, Mackrill J. Multiple Targets for Oxysterols in Their Regulation of the Immune System. Cells 2021; 10:2078.

35. Hoft SG, Sallin MA, Kauffman KD, Sakai S, Ganusov VV, Barber DL. The Rate of CD4 T Cell Entry into the Lungs during Mycobacterium tuberculosis Infection Is Determined by Partial and Opposing Effects of Multiple Chemokine Receptors. Infection and immunity 2019; 87:e00841–18.

36. Bauman DR, Bitmansour AD, McDonald JG, Thompson BM, Liang G, Russell DW. 25-Hydroxycholesterol secreted by macrophages in response to Toll-like receptor activation suppresses immunoglobulin A production. Proc Natl Acad Sci U S A 2009; 106:16764–9.

37. Park K, Scott AL. Cholesterol 25-hydroxylase production by dendritic cells and macrophages is regulated by type I interferons. Journal of leukocyte biology 2010; 88:1081–7.

38. Liu C, Yang XV, Wu J, et al. Oxysterols direct B-cell migration through EBI2. Nature 2011; 475:519–23.

39. Madenspacher JH, Morrell ED, Gowdy KM, et al. Cholesterol 25-hydroxylase promotes efferocytosis and resolution of lung inflammation. JCI Insight 2020; 5.

40. Hoff DR, Ryan GJ, Driver ER, et al. Location of Intra-and Extracellular M. tuberculosis Populations in Lungs of Mice and Guinea Pigs during Disease Progression and after Drug Treatment. PloS one 2011; 6:e17550.

41. Plumlee CR, Duffy FJ, Gern BH, et al. Ultra-low Dose Aerosol Infection of Mice with Mycobacterium tuberculosis More Closely Models Human Tuberculosis. Cell Host & Microbe 2021; 29:68–82.e5.

42. Hunter RL. Tuberculosis as a three-act play: A new paradigm for the pathogenesis of pulmonary tuberculosis. Tuberculosis 2016; 97:8–17.

43. Guillemot-Legris O, Mutemberezi V, Cani PD, Muccioli GG. Obesity is associated with changes in oxysterol metabolism and levels in mice liver, hypothalamus, adipose tissue and plasma. Sci Rep 2016; 6:19694.

44. Tremblay-Franco M, Zerbinati C, Pacelli A, et al. Effect of obesity and metabolic syndrome on plasma oxysterols and fatty acids in human. Steroids 2015; 99:287–92.

45. Fu S, Fan J, Blanco J, et al. Polysome Profiling in Liver Identifies Dynamic Regulation of Endoplasmic Reticulum Translatome by Obesity and Fasting. PLOS Genetics 2012; 8:e1002902.

