## Supplementary data for "A blunted GPR183/oxysterol axis during dysglycemia results in delayed recruitment of macrophages to the lung during *M. tuberculosis* infection"

[1]Translational Research Institute, Mater Research Institute, The University of Queensland, Brisbane, QLD 4102, Australia. [2]School of Chemistry and Molecular Biosciences, The University of Queensland, Brisbane, QLD 4072, Australia. [3]Australian Infectious Diseases Research Centre – The University of Queensland, Brisbane, QLD 4072, Australia. [4]Centre for Clinical Research, The Univeristy of Queensland, Brisbane, QLD 4072, Australia. [5]Department of Biomedical Sciences, University of Copenhagen, Copenhagen, Denmark.

* Correspondence:

Katharina Ronacher:

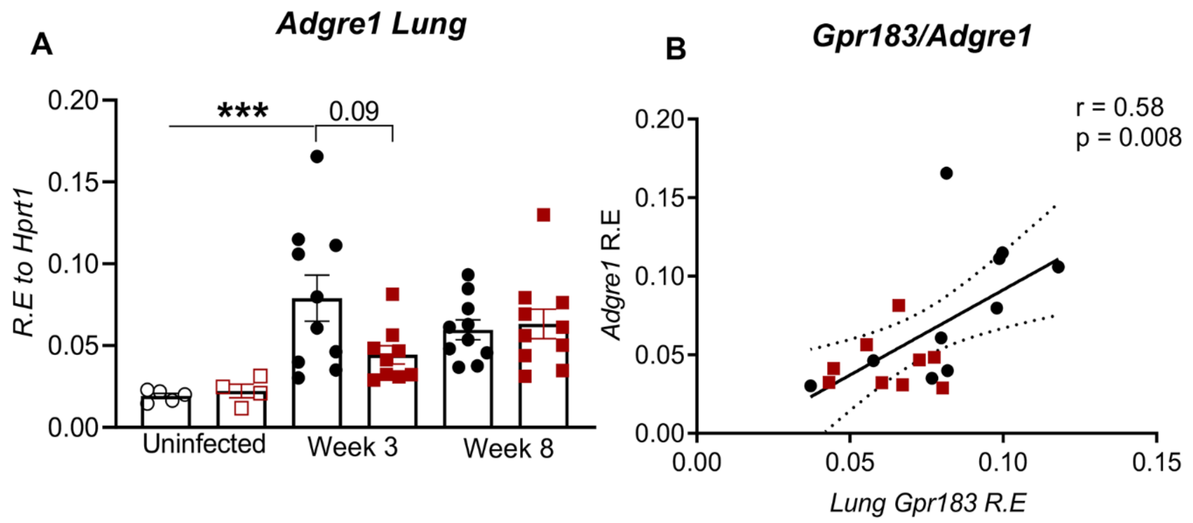


**Supplemental Figure 1: Gpr183 expression in the lung is positively correlated with the lung macrophage marker Adgre1 expression. (A)** Expression of Adgre1/F4/80 in the lung of uninfected and Mtb-infected mice with and without dysglycemia at 3 and 8 weeks p.i. **(B)**. Correlation between lung Gpr183 and Adgre1 at 3 weeks after infection using Spearman's rank correlation. (n=9-10 mice/group). Spearman r and respective p values have been shown on the figure. Circles represent normoglycemic mice and squares represent dysglycemic mice.

**Supplemental Table S1: Primers used for mRNA analysis**

| | **Forward** | **Reverse** |
|---|---|---|
| *Gpr183* | GTCGTGTTCATCCTGTGCTTCAC | TCATCAGGCACACCGTGAAGTG |
| *Ch25h* | CTGACCTTCTTCGACGTGCT | GGGAAGTCATAGCCCGAGTG |
| *Cyp7b1* | CGGAAATCTTCGATGCTCCAAAG | GCTTGTTCCGAGTCCAAAAGGC |
| *Hsd3b7* | ACTGCGCTTTGGAGGTCGTCTA | GCCACCAGTATGTGCATCCAAG |
| *Adgre1* | CGTGTTGTTGGTGGCACTGTGA | CCACATCAGTGTTCCAGGAGAC |
| *Hprt1* | CCCCAAAATGGTTAAGGTTGC | AACAAAGTCTGGCCTGTATCC |
